# Identification of a novel missense variant in *SLC45A2* associated with dilute snowdrop phenotype in Gypsy horses

**DOI:** 10.1101/851246

**Authors:** D. Bisbee, M. L. Carpenter, K. Hoefs-Martin, S. A. Brooks, C. Lafayette

**Affiliations:** Etalon Inc, Menlo Park, CA 94025, USA; Department of Animal Sciences, University of Florida, Gainesville, FL 32607, USA

## Abstract

*SLC45A2* mutations are responsible for several dilution phenotypes in horses, including *cream*, *pearl*, and *sunshine*. We sequenced the *SLC45A2* gene in a horse that possessed a diluted coat color but tested negative for any known dilution genotypes. We identified a novel homozygous missense mutation in *ECA21: SLC45A2:c.305G>A; p.(Arg102Gln)* which we have named *snowdrop* (C^sno^). The *snowdrop* dilution is autosomal recessive and dilutes both red and black pigment in the homozygous state, creating a phenotype similar to homozygous *cream*. We also identified this mutation in the heterozygous state in several relatives, including the sire and dam of the affected horse, where it has no visible effect on phenotype. Based on these data, the *snowdrop* variant produces a recessive dilution similar to *cream*, and its discovery confirms that multiple *SLC45A2* alleles cause dilute phenotypes in horses.

---

In many animals, *SLC45A2* encodes a transmembrane protein in melanocytes that is hypothesized to transport molecules necessary for proper melanosome function. Coat and skin color dilution phenotypes caused by variants in *SLC45A2* are documented in multiple vertebrate species^1–4^. In equines, mutations in the *SLC45A2* gene, located on chromosome 21, are responsible for the *cream, pearl*, and *sunshine* dilutions^5,6^.

In this study, we focused on a Gypsy breed mare that appeared phenotypically cremello (light creamy white) but tested negative for any known dilution or spotting alleles, including sabino, W20, tobiano, PATN1, and SW1-4 (Figure 1). We obtained hair samples from 15 Gypsy horses, including the affected mare, her dam and sire, and three half siblings. DNA was extracted according to a previously reported protocol^7^, and genotyping of known alleles was performed using an Agena MassArray assay. PCR and Sanger sequencing of the seven exons in the *SLC45A2* gene in the affected mare were performed as previously described^6^. Non-synonymous variants were analyzed using the PROVEAN web server^8^. Ensembl protein sequences for orthologs of *SLC45A2* in 106 mammals were downloaded from OrthoDB and aligned using clustal omega to assess the conservation of each amino acid change^9,10^.

**Figure 1.**
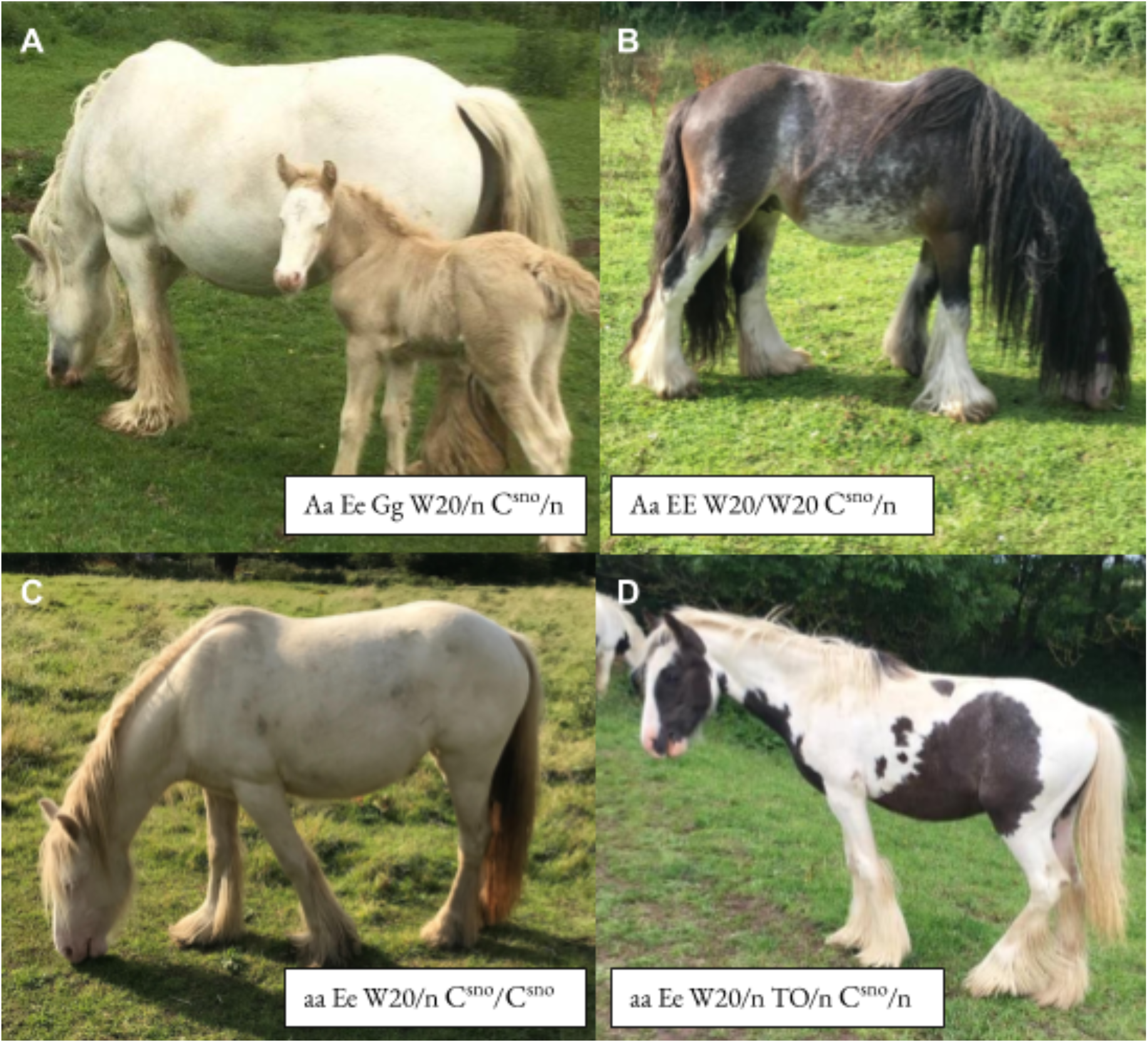
Phenotypes & genotypes of different C^sno^ positive horses. A and B are the dam and sire, respectively, of C, which is homozygous for the C^sno^ mutation. B is also the sire of D and therefore half-sibling to C.

Sanger sequencing identified three missense mutations in *SLC45A2* in the affected mare. Two variants, *SLC45A2:c.32C>A* and *SLC45A2:c.872C>A* (numbering based on transcript ENSECAT00000026240.1 in the Ensembl EquCab2.0 assembly), had positive PROVEAN scores but were in a location not conserved across vertebrates and were not investigated further. Variant *SLC45A2:c.305G>A; p.(Arg102Gln)* in exon 1 was detected in the homozygous state in the affected mare, and in the heterozygous state in the sire, dam, and a half-sibling of the mare (Table 1), and was termed *snowdrop* (C^sno^). This mutation was not identified in the other two half-siblings (Table 1) or in any of the additional nine Gypsy horses tested. Only the homozygous mare displayed a diluted phenotype. The sequence containing this variant was deposited in Genbank under accession number MN704281.

**Table 1:**
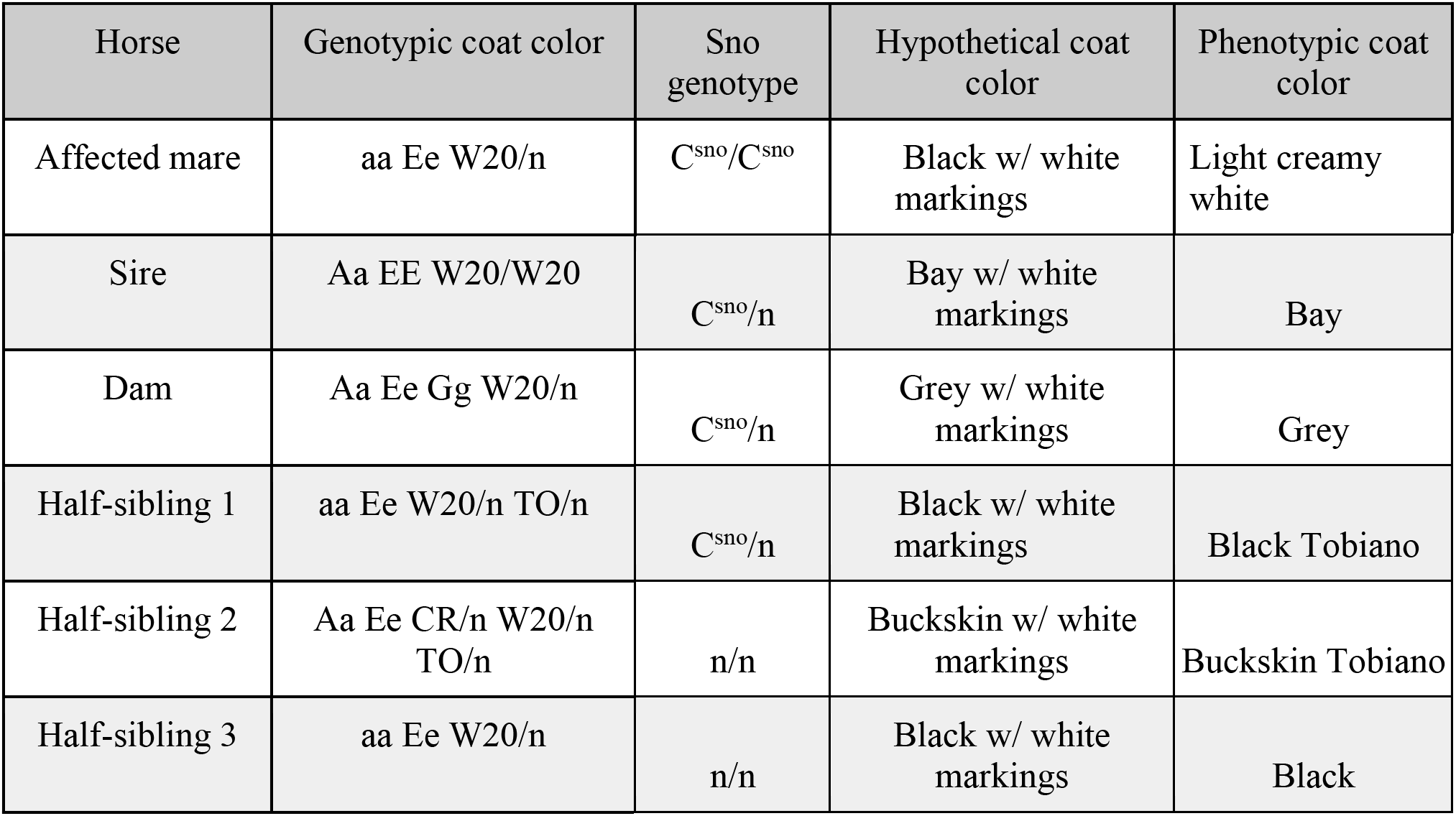
Genotypes and phenotypes of horses tested in this study.

This C^sno^ mutation at arginine 102 is predicted to be deleterious, with a PROVEAN score of −3.661. In an alignment of the *SLC45A2* protein sequence from 106 mammalian species, the horse reference allele at this position was conserved across 100% of species analyzed, also suggesting that this variant likely has an impact on protein function. The topography of the *SLC45A2* transmembrane protein is shown in Fig. S1 as predicted with PHOBIUS protein prediction software^11^; the location of the *snowdrop* allele as well as several other *SLC45A2* dilutions are shown. The C^sno^ mutation replaces the highly conserved positively charged arginine residue with a polar glutamine. Cytoplasmic regions of transmembrane proteins tend to be positively charged, a phenomenon termed the “positive-inside rule”, and disruptions in this positive charge can affect the topology and assembly of these proteins^12,13^. Thus, the *snowdrop* mutation is likely to impact the function of the *SLC45A2* protein as a membrane-associated transporter.

It is unclear whether the *snowdrop* mutation affects eumelanin or pheomelanin, but it is suspected to dilute eumelanin in the homozygous form. Homozygous *C^sno^/C^sno^* appears to dilute black pigment in both the mane and tail, and homozygous horses have pink around the nose and white eyelashes (Figure 1). Heterozygous C^sno^ appears to have no effect on phenotype; thus, C^sno^ appears to be an autosomal recessive mutation. At this time, it is unknown if the *snowdrop* dilution has any deleterious effects. Testing across additional Gypsy horses could possibly determine this new variant’s origin and allele frequency.

## Conflict of Interest

All of the authors are affiliated with Etalon Diagnostics, which offers testing of the C^sno^ variant.

## Acknowledgements

The authors would like to thank Clare Brennan, Kari Newman, and the other owners who provided samples for this study. Photos were provided by Clare Brennan.

---

**Figure S1:**
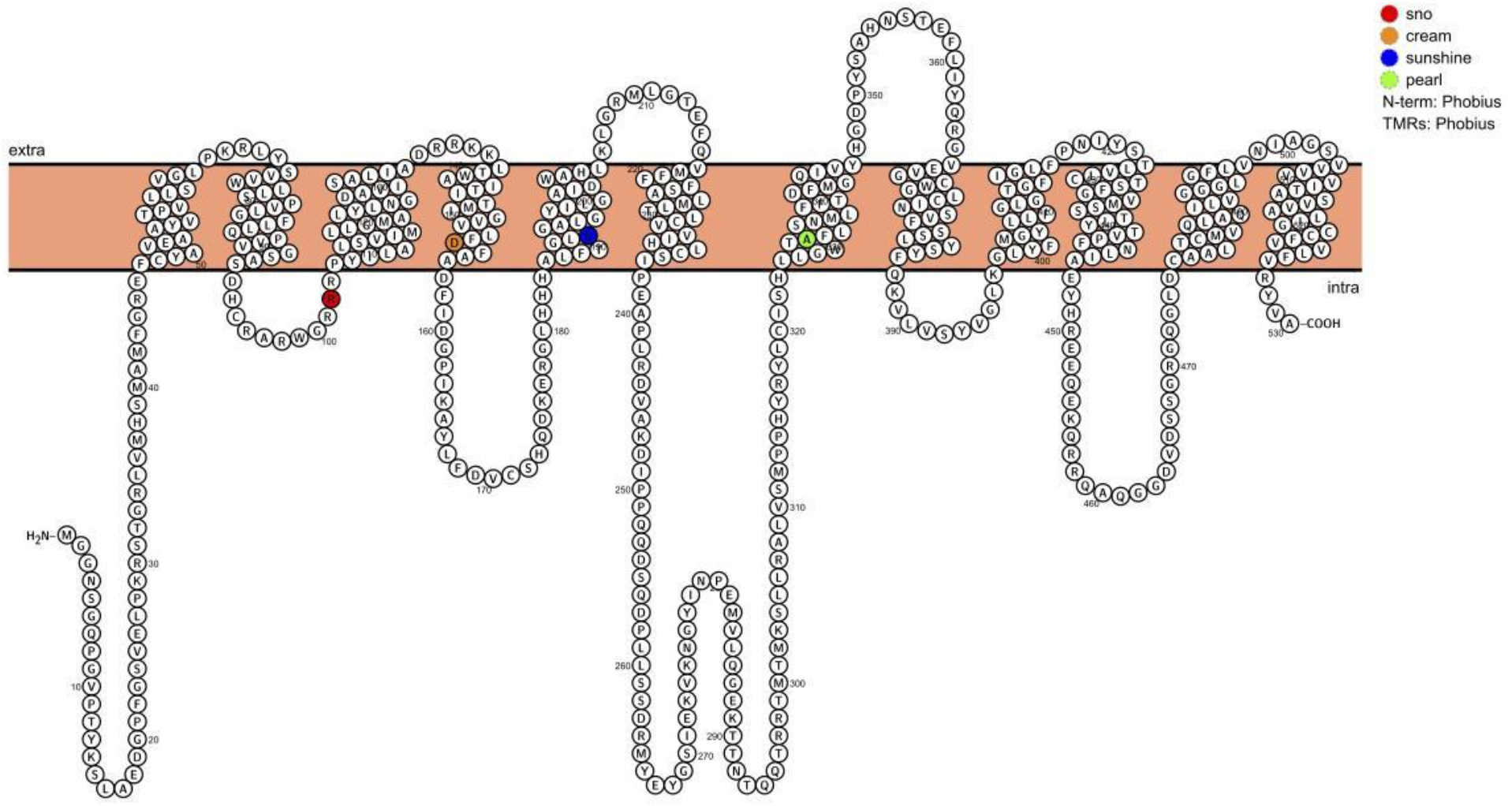
Topography of equine SLC45A2 with transmembrane regions determined using PHOBIUS protein prediction software. The *snowdrop* allele (Red) replaces a positively charged arginine residue with a neutral polar glutamine residue. The locations of previously identified mutations resulting in the *cream* (orange), *sunshine* (blue), and *pearl* (green) dilutions are shown for reference.

